# Surface area-to-volume ratio, not cellular rigidity, determines red blood cell traversal through small capillaries

**DOI:** 10.1101/2020.07.11.191494

**Authors:** Arman Namvar, Adam J. Blanch, Matthew W. Dixon, Olivia M. S. Carmo, Boyin Liu, Snigdha Tiash, Oliver Looker, Dean Andrew, Li-Jin Chan, Wai-Hong Tham, Peter V. S. Lee, Vijay Rajagopal, Leann Tilley

## Abstract

The remarkable deformability of red blood cells (RBCs) depends on the viscoelasticity of the plasma membrane and cell contents and the surface area to volume (SA:V) ratio; however, it remains unclear which of these factors is the key determinant for passage through small capillaries. We used a microfluidic device to examine the traversal of normal, stiffened, swollen, parasitised and immature RBCs. We show that dramatic stiffening of RBCs had no measurable effect on their ability to traverse small channels. By contrast, a moderate decrease in the SA:V ratio had a marked effect on the equivalent cylinder diameter that is traversable by RBCs of similar stiffness. We developed a finite element model that provides a coherent rationale for the experimental observations, based on the nonlinear mechanical behaviour of the RBC membrane skeleton. We conclude that the SA:V ratio should be given more prominence in studies of RBC pathologies.

## Introduction

The human red blood cell (RBC) comprises a plasma membrane enclosing a concentrated solution (∼10 mM) of protein, mainly haemoglobin. The mature RBC (resting long diameter ∼8 μm) exhibits remarkable deformability and durability. During its four-month lifespan, a human RBC circulates the body about a million times (Allison, 1960). It undertakes this journey of about 500 hundred kilometres, without repair, squeezing through cerebral capillaries with diameters as small as 3 μm and inter-endothelial slits in the spleen with widths of 1-2 μm (An *et al*., 2008, Lauwers *et al*., 2008, Marin-Padilla, 2012). The ability to undergo large deformations and to conform to narrow shapes is essential for RBC function.

The deformability of RBCs is determined by contributions from the plasma membrane viscoelasticity, the cytoplasmic viscosity and the cellular geometry (Mohandas *et al*., 2008, Diez-Silva *et al*., 2010, Pivkin *et al*., 2016). The viscoelastic properties of the RBC membrane are in turn determined by a sub-membranous protein meshwork (Mohandas *et al*., 2008, Zhang *et al*., 2015). The membrane skeleton comprises an array of flexible cross-members (spectrin) linked into junctional complexes (actin and accessory proteins) (Mankelow *et al*., 2012). The spectrin molecules act like molecular springs, accommodating the distortions imposed by shear forces in the circulation. Vertical interactions connect the skeletal meshwork to the proteins embedded in the plasma membrane. The cytoplasmic viscosity is dependent on the concentration and physical state of haemoglobin, which may be altered by changes in the hydration level or in some haemoglobinopathies (Kuhn *et al*., 2017, Renoux *et al*., 2019). A resting RBC adopts a biconcave discoid shape that optimises the surface area to volume (SA:V) ratio (∼1.5-fold greater than a sphere of the same volume). The surface area of a healthy RBC is kept constant, and any increases in volume decrease the SA:V ratio, which adversely impacts RBC rheology.

Upon infection with a malaria parasite, key biomechanical properties of the RBC are subverted. For example, RBCs infected with mature stage *Plasmodium falciparum* - the species responsible for most malaria-related deaths - exhibit significantly decreased deformability (Glenister *et al*., 2002, Suresh *et al*., 2005, Park *et al*., 2008). The intracellular parasite exports proteins that stiffen the RBC membrane (de Koning-Ward *et al*., 2016). *P. falciparum* also exports an adhesin, called *P. falciparum* Erythrocyte Membrane Protein-1, that embeds in the RBC membrane and binds to endothelial cell receptors (Wahlgren *et al*., 2017). Sequestration of infected RBCs in brain capillaries is associated with impaired blood flow within the deep tissue microcapillary beds (Warrell *et al*., 1988, Dondorp *et al*., 2000, Beare *et al*., 2009). In some cases this leads to a serious complication, known as cerebral malaria, which is associated with coma and neurological sequalae (Renia *et al*., 2012). It remains unclear to what extent the higher rigidity of infected RBCs contributes directly to trapping of infected RBCs.

*P. knowlesi* is a parasite of long-tailed macaques (*Macaca fascicularis*) that causes zoonotic infections in humans and is the most common cause of human malaria in Malaysia (Singh *et al*., 2013). It is associated with high parasitemia infections and causes severe malaria in adult humans at a rate similar to *P. falciparum*(Cox-Singh *et al*., 2008, Barber *et al*., 2013). *P. knowlesi* does not express adhesins on the surface of the host RBC, but does exhibit increased the host cellular rigidity(Barber *et al*., 2018b). Despite the lack of adhesion, splenic rupture has been observed, suggesting blockage of sinusoidal vessels(Chang *et al*., 2018) and an autopsy of a fatal human knowlesi malaria case revealed brain capillaries congested with infected RBCs(Menezes *et al*., 2012). Acute kidney injury, potentially caused by chronic haemolysis, also contributes to severe pathology(Barber *et al*., 2018a). The molecular basis for the different pathologies remains poorly understood(Renia *et al*., 2012).

Altered cellular properties are also observed in immature RBCs, known as reticulocytes. The earliest enucleated reticulocyte forms are released from the bone marrow with extra surface area and an elevated membrane shear modulus(Liu *et al*., 2010, Malleret *et al*., 2013). The transition of reticulocytes to mature RBCs is accompanied by the loss of surface area, acquisition of a biconcave shape and increased deformability(Li *et al*., 2018). The consequences of the lower deformability of early stage reticulocytes on passage through small capillaries is not well studied.

RBC deformability is a major determinant of the behaviour of blood cells in the circulation. A technique called ektacytometry is routinely employed as one measure of deformability. Ektacytometry measures the ability of RBCs to elongate upon application of shear stress in fluid flow. Membrane stiffness and cytoplasmic viscosity are important determinants of the ability of RBCs to elongate(Mohandas *et al*., 2008, Diez-Silva *et al*., 2010, Pivkin *et al*., 2016), while the contribution of SA:V ratio is less well studied. Another physiologically relevant parameter is the ability of RBCs to passage into small constrictions, which is expected to have consequences for transfer through very narrow capillaries and splenic fenestrations(Kim *et al*., 2012, Safeukui *et al*., 2018). However, the individual effects of cell geometry and cellular rigidity on capillary traversal are not well understood.

In this study, we monitored RBC traversal into an array of microchannels, with diameters commensurate with the smallest vessels in the circulation and examined the effects of independently manipulating cellular rigidity and SA:V ratio. As examples of physiological and pathological changes to RBC membrane properties, we examined the traversal of reticulocytes and *P. falciparum-* and *P. knowlesi*-infected RBCs. We developed a computational model and used it to help understand the contributions from different parameters and estimate the minimum pressure required to force RBCs with different SA:V ratios and cell stiffness properties through small capillaries, mimicking the dimension of those in the cerebral cortex.

## Results

### Calibrating measurements of RBC geometry

We employed a modified version of the microfluidic device called a Human Erythrocyte Microchannel Analyser (HEMA) to assess the ability of RBCs to traverse into wedge-shaped microchannels. Scanning electron microscopy (SEM) images of a chip are presented in Supplementary Fig. 1. RBCs enter the ∼5 μm diameter opening and become trapped before reaching the 1.4 μm exit (Fig. 1a). The position where the RBC becomes lodged represents the smallest diameter of an equivalent cylindrical tube through which the RBC can pass. We use a previously coined term, minimum cylindrical diameter (MCD)(Canham *et al*., 1968, Herricks *et al*., 2009) to describe this parameter. Subtle differences in the way MCD is calculated in our study are detailed in the Methods section. Conforming the RBCs into the microchannels allows estimation of the cell surface area and volume(Gifford *et al*., 2003, Herricks *et al*., 2009, Lelliott *et al*., 2017).

**Figure 1.**
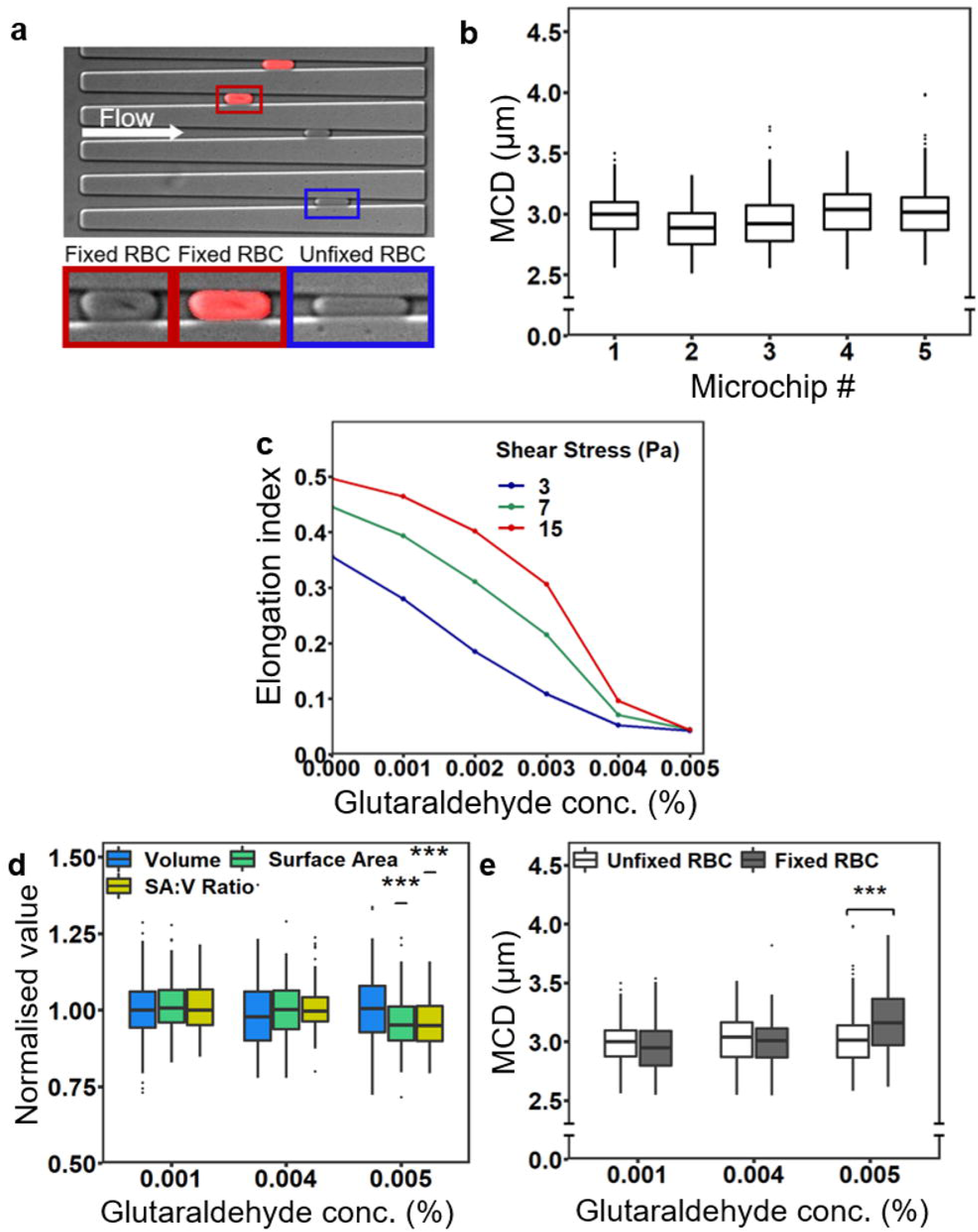
Ektacytometry and HEMA analyses of unfixed and fixed RBCs. (a) An image of the wedge-shaped microchannels in the HEMA chip with arrested unfixed RBCs (blue box) and 0.005% glutaraldehyde-fixed RBCs (red autofluorescence, red box), with one cell showing a crease (see inset with no fluorescence overly). (b) Comparison of the minimum cylindrical diameter (MCD) for healthy RBCs examined in different HEMA chips. (c) Comparison of the elongation index (EI) for RBCs treated with glutaraldehyde, at shear stresses of 3, 7 and 15 Pa. (d) Normalised values of SA:V ratio for fixed and unfixed RBCs. Individual data points indicate the full range of values. Data represent a single experiment. An additional experiment is presented in Supplementary Fig. 2b,c. (e) Mean MCD values ± S.D. for unfixed RBCs (white) and fixed RBCs (grey).

It was important to determine the variability in the measurements, which could be affected by batch-to-batch differences in microchannel dimensions. We analysed the same batch of RBCs in five independently fabricated chips, revealing small variations in estimates of MCD (Fig. 1b). In an effort to obtain a reliable estimate of the different geometrical parameters, we averaged data from eight different experiments (2562 RBCs). This analysis reveals an average volume of 99 ± 12 fL and an average surface area of 149 ± 14 µm^2^, giving a SA:V ratio of 1.50 ± 0.12. The average MCD is 3.03 ± 0.21 µm. The data are in good agreement with previous reports(Gifford *et al*., 2003, Park *et al*., 2008, Herricks *et al*., 2009, Hanssen *et al*., 2012). Nonetheless, given the degree of variation, where possible we examined different samples in the same chip by differentially labelling one population or washing and reusing the chip for different samples on the same day. In some experiments, the data was normalised to the untreated RBCs.

### Cell stiffness limits RBC elongation but not traversal into microchannels

We first examined the behaviour of RBCs under conditions in which the cellular rigidity was modified by chemical treatment, but the SA:V ratio was held as constant as possible. Glutaraldehyde is a non-specific cross-linking agent that stiffens the cell membrane by cross-linking membrane proteins, and increases cellular viscosity by cross-linking haemoglobin(Forsyth *et al*., 2010). RBCs were treated for 1 h with different glutaraldehyde concentrations. Here, we refer to the composite change in properties as an increase in cellular rigidity.

We used a RheoScan ektacytometer(Shin *et al*., 2005) to measure the elongation index (EI) of RBCs at shear stresses ranging from 0-20 Pa. RBCs were mixed with 6% polyvinylpyrrolidone (PVP) in buffer prior to analysis. Ektacytometry revealed a marked decrease in EI as the glutaraldehyde concentration increases from 0.001% to 0.005% (Fig. 1c). At the physiologically relevant shear stress of 3 Pa, the ability of the RBCs treated with 0.004% glutaraldehyde to elongate was largely abrogated (Fig. 1c).

For HEMA analysis, glutaraldehyde-treated RBCs and control (unfixed) RBC were mixed in phosphate-buffered saline (PBS) at a ratio of 1:1 and introduced into the microfluidic chip. Fixed RBCs were identified by an increased autofluorescence signal (Fig. 1a). Different chips were used for the three glutaraldehyde concentrations examined and the data were normalised to the untreated RBCs in the same chip.

No significant changes in volume or surface area, and thus SA:V ratio, were observed between fixed and unfixed RBCs at glutaraldehyde concentrations of 0.001 and 0.004% (Fig. 1d). (The actual values for the SA:V ratio are provided in Supp Fig. S2a). These data show that markedly stiffened RBCs (0.004% glutaraldehyde) retain the ability to conform to the shape of the narrowing microchannel, reaching an MCD of 3.00 ± 0.23 µm, similar to unfixed RBCs (3.02 ± 0.21 µm; P = 0.54; 95% confidence interval: -0.075 to 0.039; n = 279 unfixed RBCs, 76 fixed RBCs). These data show that cellular rigidity has little impact on RBC traversal into small capillaries.

Treatment with 0.005% glutaraldehyde increases the MCD to 3.19 ± 0.31 µm, which is significantly larger than for unfixed RBCs measured in the same chip (3.02 ± 0.22 µm) (P < 0.00001; 95% confidence interval: 0.11 to 0.24; n = 257 unfixed RBCs, 104 fixed RBCs; Fig. 1d). In this case, however, the surface area, and thus SA:V ratio, appears to be decreased compared with unfixed RBCs (Fig. 1e; Supplementary Fig. 2). This difference emanates from the formation of creases in the membrane (see Fig. 1a), causing an underestimation of the surface area. No creases were observed in the membranes of RBCs fixed with lower glutaraldehyde concentrations or in unfixed RBCs (see Supplementary Fig. 3).

### SA:V ratio limits traversal of RBCs into microchannels but has less effect on elongation

We next examined the impact of altered cell volume on microchannel traversal and RBC elongation, by varying the osmolarity of the buffering solution from 139-484 mOsm/L. At higher or lower osmolarities, the ability of RBCs to elongate in flow decreased substantially (Fig. 2a), in agreement with previous data(Clark *et al*., 1983, Renoux *et al*., 2019). However, at osmolarities relatively close to the physiological range, 239-350 mOsm/L, the EI showed relatively little dependence on osmolarity (Fig. 2a, red box). We examined RBC behaviour over the same range, using a single HEMA device, with flushing of the device between experiments. We observed a gradual decrease in cell volume and a slight apparent increase in cell surface area as the cells dehydrate and shrink in the increasing buffer osmolarity (Fig. 2b,c; Supplementary Fig. 4). The resultant increase in SA:V ratio (from 1.38 ± 0.11 to 1.58 ± 0.10; Fig. 2d) is associated with a decrease in the MCD value from 3.29 ± 0.24 µm (239 mOsm/L; n = 411 RBCs) to 2.89 ± 0.15 µm (350 mOsm/L; n = 423 RBCs; p < 0.00001; 95% confidence interval: 0.37 to 0.43; Fig. 2e). Indeed, a roughly inverse linear relationship was observed between MCD and SA:V ratio (Supplementary Fig. 4).

**Figure 2.**
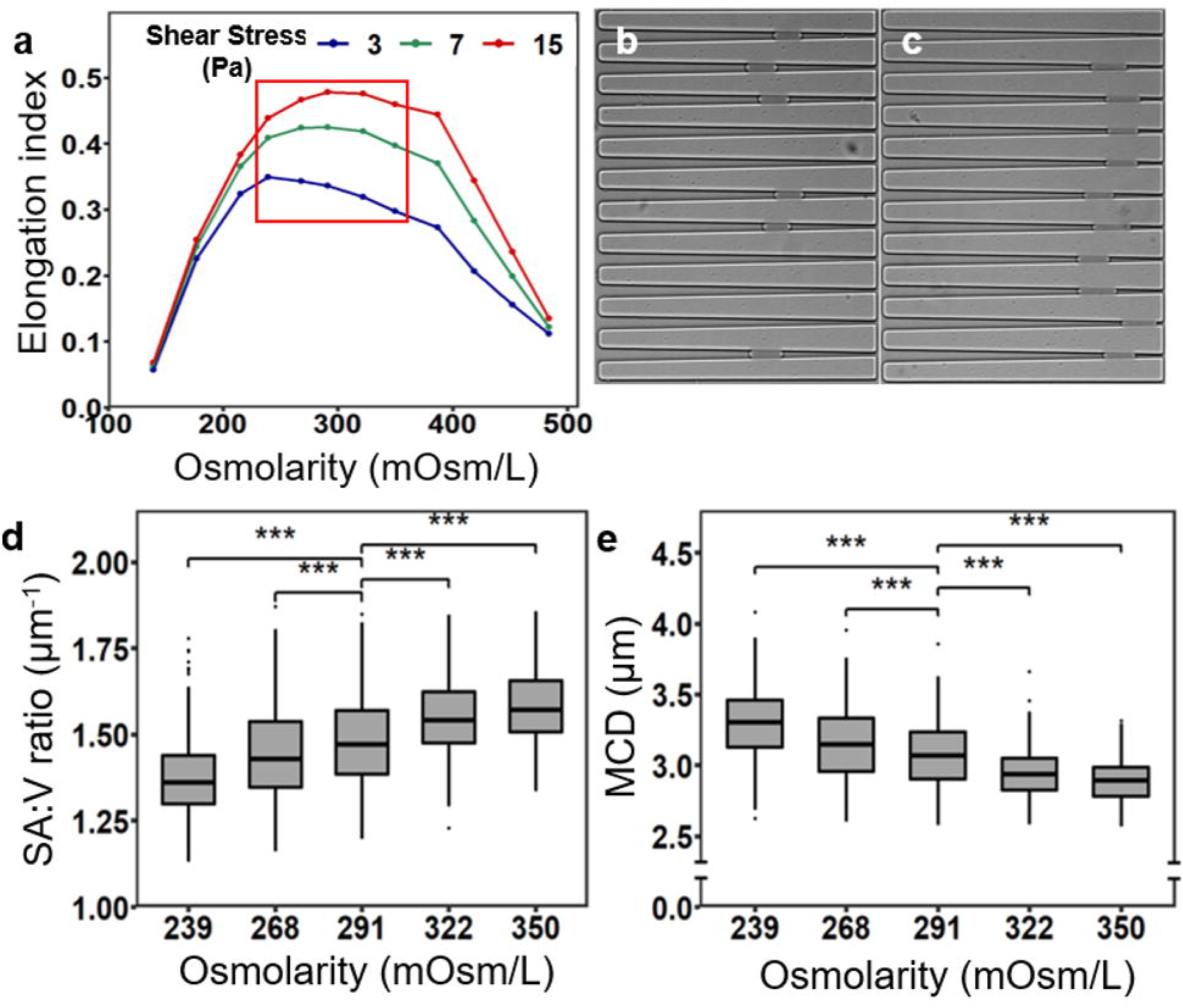
Ektacytometry and HEMA analyses of RBCs subjected to different osmolarities. (a) Elongation index as a function of the medium osmolarity at shear stresses of 3, 7 and 15 Pa. (b-c) Images of the wedge-shaped microchannels with arrested RBCs in media with osmolarities of 239 mOsm/L (b) and 350 mOsm/L (c). (d,e) Mean values SA:V ratio and MCD for RBCs subjected to different osmolarities. All five conditions were examined in a single HEMA chip, starting from 239 mOsm/L. After each experiment, the chip was flushed with the next buffer for 30 min before re-introducing the RBCs. The data are from a single experiment and are typical of data obtained in five separate experiments.

### SA:V ratio is the main determinant of microchannel traversal for malaria parasite-infected RBCs

We next examined the importance of SA:V ratio and cell stiffness in determining microchannel traversal of parasitised RBCs. RBCs infected with *P. falciparum* (3D7) trophozoites (32-36 h) were magnet-purified and washed in complete (serum-containing) culture medium and analysed by ektacytometry at a parasitemia level of 98%. The infected RBCs exhibit markedly increased cellular rigidity compared with uninfected RBCs (Fig. 3a). *P. falciparum*-infected RBCs exhibited a rigidity profile similar to that of RBCs treated with 0.005% glutaraldehyde.

**Figure 3.**
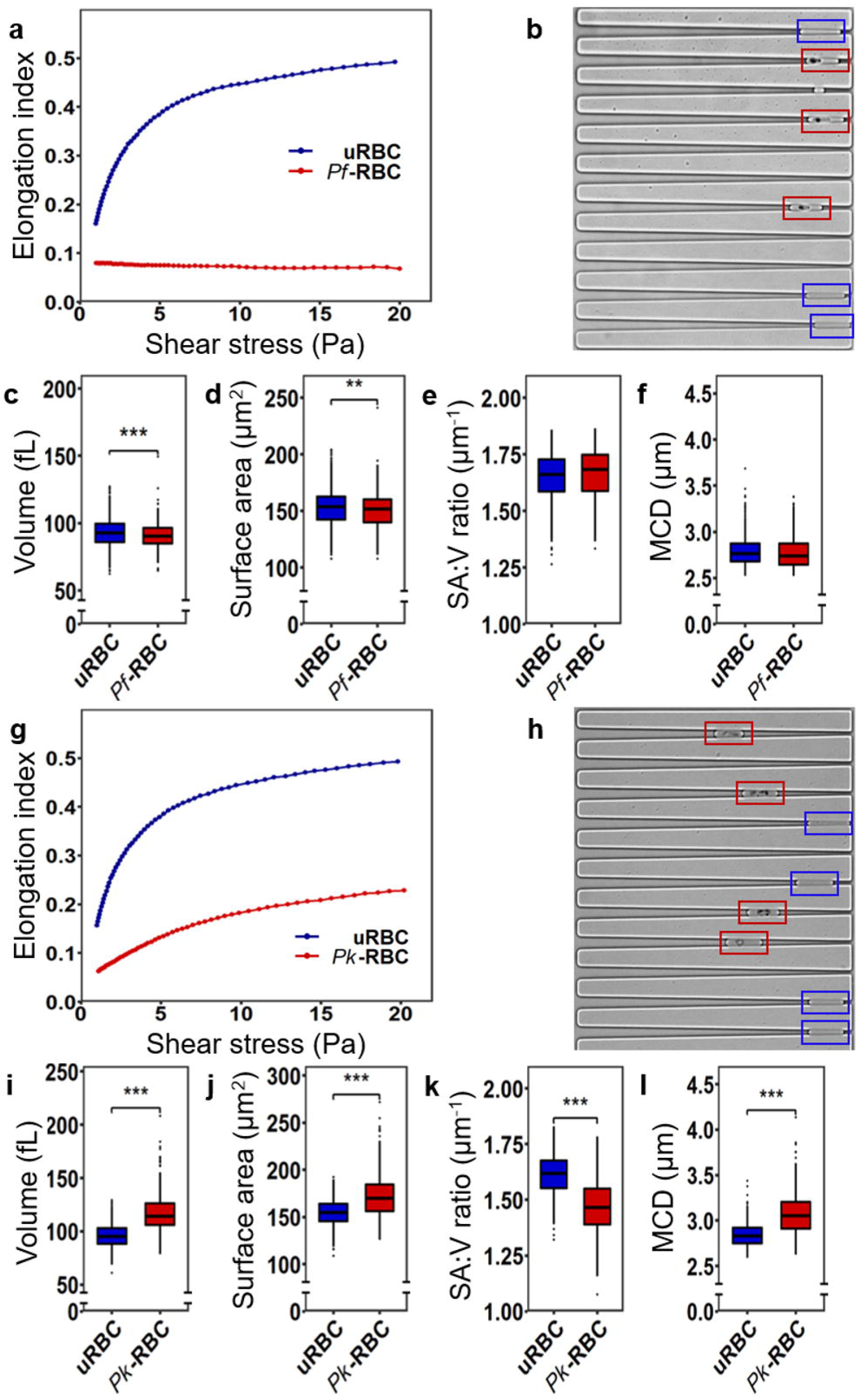
Ektacytometry and HEMA analyses of parasitised RBCs. (a,g) Ektacytometry (0-20 Pa shear stress) of uninfected RBCs (blue line) and RBCs infected with *P. falciparum* (a; 32-36 h) (red line) or *P. knowlesi* (g; 22-26 h). (b,h) RBCs infected with *P. falciparum* (b) or *P. knowlesi* (h) (red boxes) and uninfected RBCs (blue boxes) were trapped in wedge-shaped microchannel. Mean values of volume, surface area, SA:V ratio and MCD for RBCs infected with *P. falciparum* (c-f, red) or *P. knowlesi* (i-l) and uninfected RBCs (blue). The data represent combined results from five experiments for *P. falciparum* and three experiments for *P. knowlesi*.

For the HEMA experiments we directly compared uninfected and infected RBCs in the same chip. We used the highly absorbing hemozoin crystals to identify infected RBCs (Fig. 3b). It should be noted that the MCD for uninfected RBCs in complete culture medium (MCD = 2.81 ± 0.15; n = 1528 RBCs) was moderately lower than for RBCs suspended in PBS (MCD = 3.03 ± 0.21; n = 2562 RBCs) (P < 0.00001; 95% confidence interval: -0.23 to -0.21), potentially due to nutrient or serum albumin effects.

We observed no significant difference in the mean SA:V ratio of *P. falciparum* trophozoites compared to uninfected RBCs, measured in the same chip (Fig. 3c-f), in agreement with some previous reports(Herricks *et al*., 2009, Hanssen *et al*., 2012, Liu *et al*., 2019). Despite the very limited ability of *P. falciparum* trophozoite-infected RBCs to elongate in fluid flow and the presence of a rigid parasite inside the host cell, infected RBCs were able to traverse to the same MCD value (2.78 ± 0.18; n = 529) as uninfected RBCs (2.79 ± 0.15; n = 1038; p = 0.18; 95% confidence interval: -0.029 to 0.0056; five experiments; Fig. 3f). The same behaviour was observed for RBCs infected with trophozoites of another strain of *P. falciparum* (CS2; Supplementary Fig. 5).

We also examined RBCs infected with *P. knowlesi* trophozoites (22-26 h). Magnet-purified infected RBCs (in serum-containing culture medium) were analysed by ektacytometry at a parasitemia level of 97%. *P. knowlesi*-infected RBCs show reduced ability to elongate in flow (Fig 3g), but appeared less rigid than *P. falciparum*-infected RBCs. *P. knowlesi* trophozoite-infected RBCs exhibit an increased volume (∼23%) and surface area (∼12%) compared to uninfected RBCs, leading to a ∼9% decrease in the SA:V ratio (Fig 3i-k). Accordingly, the *P. knowlesi*-infected RBCs (MCD 3.08 ± 0.23; n = 406 RBCs) were less able to traverse microchannels than uninfected RBCs (MCD 2.84 ± 0.13; n = 490 RBCs) (P < 0.00001; 95% confidence interval: 0.21 to 0.26; three experiments; Fig. 3h,l).These data confirm that SA:V ratio is a more important determinant than cell stiffness of the ability of RBCs to traverse small microchannels, with implications for the pathology of *P. knowlesi* infections.

### Reticulocytes have an equivalent SA:V ratio to mature RBCs and traverse microchannels with similar efficiency despite higher stiffness

Immature RBCs, known as reticulocytes, have a higher surface area and a higher shear modulus than mature RBCs(Malleret *et al*., 2013). Here we purified reticulocytes by centrifugation on a Percoll cushion. Block-face SEM revealed the larger size and lobed nature of the reticulocytes (Supplementary Fig. 6; see Supplementary Video 1 for rotations of the models). The reticulocyte fraction contains a variable population of more mature RBCs (negative for the Thiazole Orange RNA stain and/or CD71 (*i*.*e*. transferrin receptor)). The EI at 3 Pa of reticulocyte-enriched samples (80% RNA positive) was approximately 13% lower than for mature RBCs (n=3, representative plot in Fig. 4a), consistent with previous reports of a two-fold higher shear modulus, as measure by micropipette aspiration(Malleret *et al*., 2013).

The reticulocyte fraction was labelled with an antibody recognising CD71, which is lost during maturation(Kono *et al*., 2009). Mature RBCs retrieved from the Percoll pellet fraction were counter-labelled with an antiserum recognising complement receptor-1 (CR1), which is present on all maturation stages. The reticulocyte faction (CD71+ and CD71-) and mature RBCs were mixed and introduced into the HEMA device in PBS (Fig. 4b). The volume and surface area of CD71+ reticulocytes were each found to be ∼30% higher than for mature RBCs (Fig. 4c,d), while CD71-reticulocytes exhibited intermediate sizes. Despite the difference in size, both CD71+ and CD71-reticulocytes exhibit the same SA:V ratio as mature RBCs (Fig. 4e) and were found to reach the same MCD (Fig. 4f; CD71+ reticulocyte MCD = 3.09 ± 0.18, n = 168; mature RBC MCD = 3.10 ± 0.15, n = 69; p value = 0.79; 95% confidence interval: -0.052 to 0.039).

**Figure 4.**
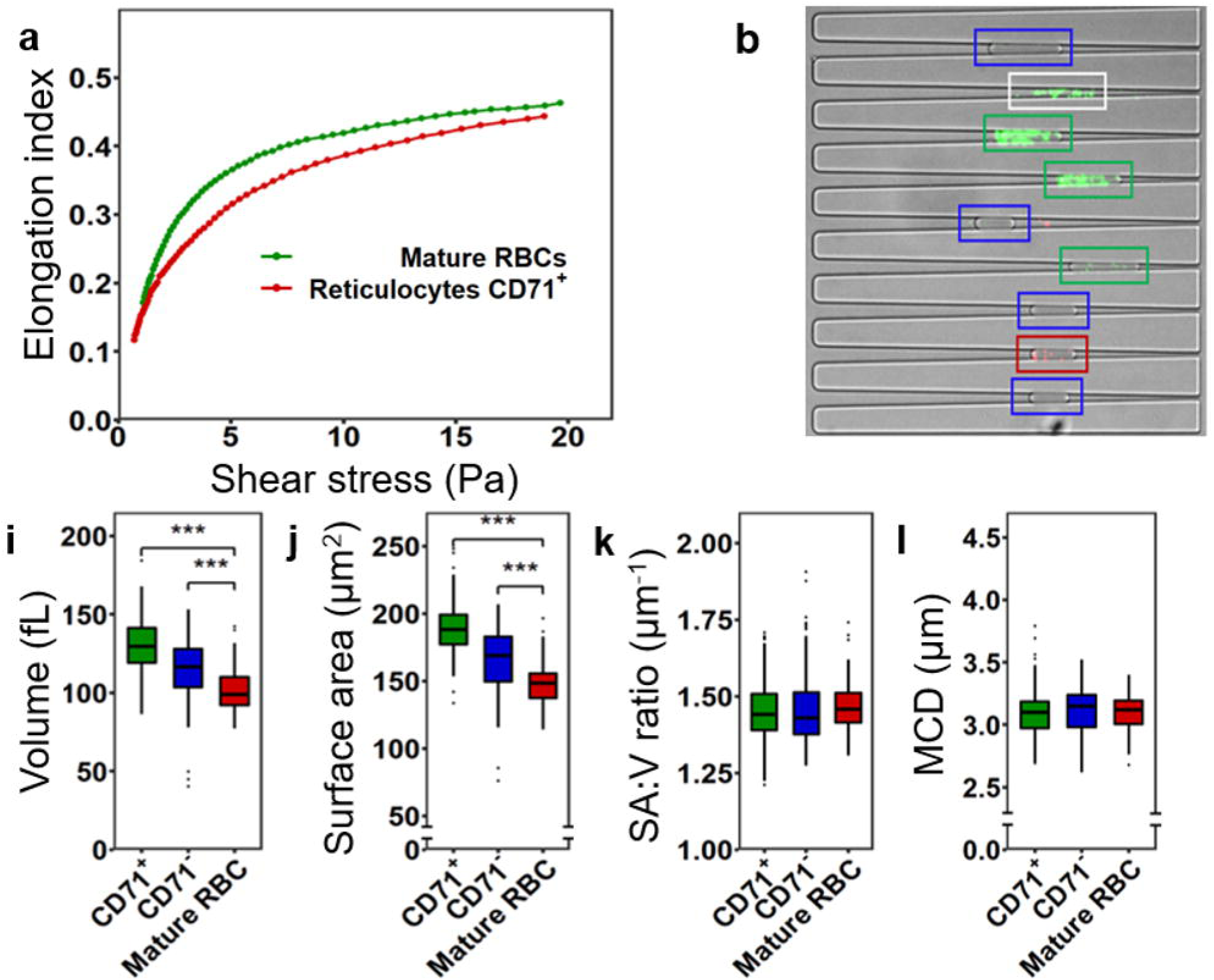
Ektacytometry and HEMA analyses of reticulocytes. (a) Ektacytometry (0-20 Pa shear stress) of uninfected RBCs (red line) and purified cord blood reticulocytes (80% RNA positive, 59% CD71+; green line). (b) Mature RBCs were labelled for CR1 (647, red boxes) and reticulocytes (purified from whole blood; 66% RNA+ positive, 33% CD71+) for CD71 (488, green boxes). Unlabelled cells are assumed to be CD71-cells from the reticulocyte fraction (blue boxes). Some lysed cell membranes are visible in the channels (white boxes). (e-f) Comparison of mean values of volume (c), surface area (d), SA:V ratio (e) and MCD (f) for mature RBCs (red), intermediate stages (CD71-, blue) and reticulocytes (CD71+, green). The data are from a single experiment and are typical of data obtained in three separate experiments.

### A 3D computational model of RBCs traversing the HEMA microchannels

We developed a biomechanical simulation model of RBC traversal into the HEMA chip to estimate the forces that RBCs would experience, and to probe the physical basis for the importance of SA:V ratio on RBC traversal through narrow microchannels. We deployed the commercial Abaqus software (SIMULIA, Providence, RI, USA) to generate a nonlinear 3D Finite Element (FE) model (Fig. 5a), building on previous studies(Mills *et al*., 2004, Aingaran *et al*., 2012). The membrane was modelled using the third order Yeoh hyper-elastic model(Yeoh, 1990) with an initial in-plane shear modulus of μ_0_ = 7.3 μN/m for healthy RBCs, based on reported values(Mills *et al*., 2004, Yoon *et al*., 2016). We employed an initial SA:V ratio value of 1.54, based on our own initial estimate of SA:V ratio. For consistency with previously generated models(Evans *et al*., 1972), we also examined a SA:V ratio of 1.42 (Supplementary Fig. 7, 9). Further details of the model can be found in the Methods section.

**Figure 5.**
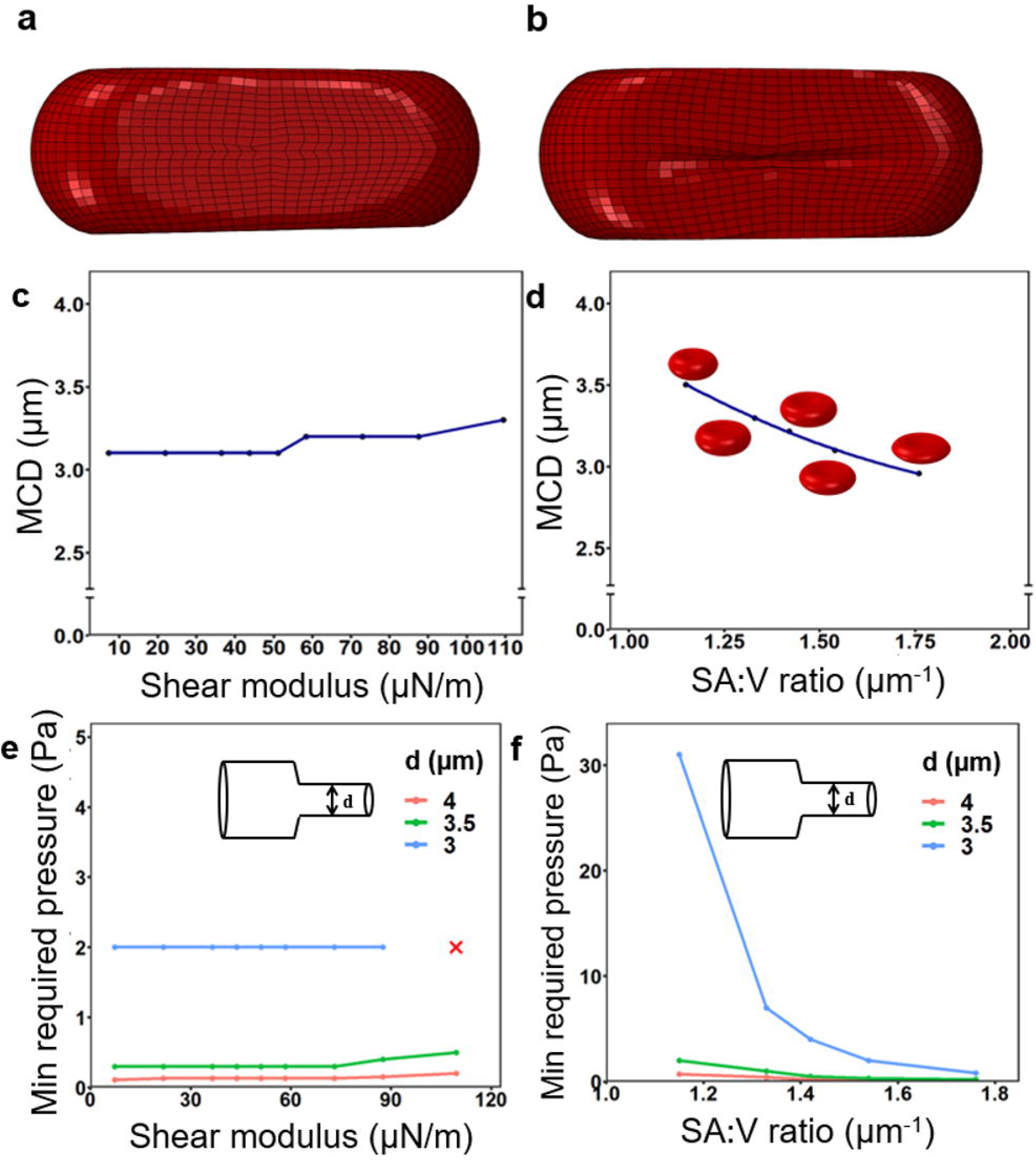
Numerical simulations of RBC passage through microchannels and capillaries. (a,b) Deformed shapes of RBCs subjected to μ = 7.3 μN/m (a) and μ = 73 μN/m (b), showing strain around creases in the membrane. (c) MCD reached by RBCs in the microchannels as a function of increasing membrane shear modulus. (d) MCD reached by RBCs at SA:V ratios of 1.15 to 1.76. (e,f) Simulations of RBC traversal in small capillaries (3 to 4 μm). The minimum required pressure is predicted for RBCs with different membrane shear moduli (e) and SA:V ratios (f).

At any given point in the HEMA device, an RBC undergoes a complex distribution of stresses that cannot be measured experimentally. Thus, we resorted to estimating the pressure that an RBC would experience using our FE model and HEMA measurements of deformed length and diameter of healthy RBCs. An RBC is subjected to a net force along the flow direction when it is introduced to the microchip due to the pressure difference (drop) across the cell surface. The pressure drop per cell increases as the RBC squeezes into the narrowing microchannel, and the pressure-drop reaches its maximum once the cell is trapped. To estimate the pressure-drop per RBC in the wedge-shaped channels, we applied a range of pressures to the HEMA simulation and found that a pressure drop of 2.20 Pa per cell best fits our experimental measurements on the untreated RBCs (Supplementary Fig. 7). The estimated pressure drop value is similar to that reported in previous studies(Aingaran *et al*., 2012, Pivkin *et al*., 2016). All subsequent simulations of the HEMA experiments on chemically treated RBCs were conducted with the estimated pressure drop of 2.20 Pa.

### Simulation of the effect of membrane stiffness on RBC traversal into a microchannel

We carried out two numerical analyses of RBCs within the HEMA device, considering (i) only membrane stiffness variation, and (ii) only SA:V ratio variation. The in-plane membrane shear modulus is reported to be in the range of 5-10 μN/m for normal RBCs(Evans *et al*., 1984, Hochmuth *et al*., 1987), increasing to ∼40 µN/m for trophozoite stage *P. falciparum*-infected RBCs, and to as high as 90 µN/m at the schizont stage(Suresh *et al*., 2005, Park *et al*., 2008). RBCs infected with mature stage *P. knowlesi* exhibit a membrane shear modulus of 45 μN/m(Barber *et al*., 2018b). We simulated variation of in-plane membrane shear modulus with values from 7.3 to 109.5 μN/m and examined the MCD reached for RBCs with a constant SA:V ratio of 1.54 (Fig. 5c). RBCs exhibiting a shear modulus up to 50 μN/m were trapped in the microchannels at an MCD of 3.1 μm (Fig. 5c), which is similar to the experimentally determined value for RBCs with the equivalent SA:V ratio (Fig. 2e). RBCs exhibiting a shear modulus greater than 50 μN/m exhibit a moderate decrease in traversal into the microchannels. For instance, an RBC exhibiting a shear modulus of 109.5 μN/m exhibited a ∼6% larger MCD than a healthy cell. Creases formed in the cell membrane in the simulations when the membrane shear modulus was higher than 73 μN/m (Fig. 5b), similar to those observed experimentally in RBCs treated with 0.005% glutaraldehyde. The simulation is consistent with the observation that a dramatic increase in the membrane shear modulus has only a moderate effect on RBC traversal into the microchannels. The range of stresses and strains in the membrane at the various shear moduli is presented in Supplementary Fig. 8. We also examined the effect of membrane stiffness on the passage of RBCs with a SA:V ratio of 1.42 (Supplementary Fig. 9).

### Simulation of the effect of SA:V ratio on RBC traversal into a microchannel

Simulations were performed on RBCs with different SA:V ratios, ranging from 1.15-1.76. These RBC models were generated by altering the geometrical parameters in the standard geometry formula(Evans *et al*., 1972). The volumes of the generated cells were varied from 117 to 78 fL, while keeping the surface area constant (134 μm^2^) and the membrane shear modulus constant (μ_0_ = 7.3 μN/m). The MCD decreased as the SA:V ratio increased (Fig. 5d), consistent with the observation that the passage of RBCs into the microchannels is highly dependent on the SA:V ratio. The range of stresses and strains in the membrane at the different SA:V ratios is detailed in Supplementary Fig. 8.

### Prediction of RBC passage in cerebral capillaries

We used our whole cell FE model to predict whether high cell stiffness or low SA:V ratio would restrict the passage of a population of RBCs through the smallest cerebral capillaries encountered under patho/physiological conditions, which may have diameters as low as 3 μm(Canham *et al*., 1968, Lauwers *et al*., 2008, Linninger *et al*., 2013). In this work, we found that the average MCD for healthy RBCs suspended in PBS is 3.03 ± 0.21 μm (average of ∼2562 RBCs from different experiments). It should be noted, however, that the 90 percentile values for MCD are 2.73 and 3.29 μm. We performed simulations to evaluate the minimum required pressure for RBCs with a SA:V ratio of 1.54 to squeeze into small capillaries with diameters of 3, 3.5 and 4 μm (Fig. 5e,f). RBCs pass through the 3.5 and 4 μm capillaries, even under very low pressures; however, a higher pressure (∼2 Pa) was required to drive RBCs through the 3 μm channel. Increased cellular rigidity did not affect the minimum required pressure for RBCs to pass through these small capillaries (Fig. 5e). All stiffened cells except the one with the highest shear modulus (109.5 μN/m) can pass the minimum diameter capillary (3 μm) by applying a pressure drop of 2 Pa. We also examined the minimum required pressure for RBCs with a SA:V ratio of 1.42 and different levels of stiffness (Supplementary Fig. 9). Again, shear modulus is not an important determinant.

In contrast, the minimum required pressure steeply increases as the SA:V ratio decreases, most strikingly for the 3 μm channel. For RBCs with a SA:V ratio less than 1.4, a marked increase in the pressure is required to drive traversal through the 3 μm capillary (Fig. 5f). Our simulations show that RBCs with a low SA:V ratio are more prone to trapping in small cerebral capillaries than cells with high cellular rigidity.

## Discussion

RBCs need to undergo severe deformation to pass through narrow capillaries and fenestrations; and their ability to do so can be compromised in different physiological and pathological conditions(Suzuki *et al*., 1996, Tomaiuolo, 2014, Safeukui *et al*., 2018). Over the past few decades, many studies have examined the biomechanical properties of healthy and diseased RBCs using a variety of tools. Techniques such as micropipette aspiration, Atomic Force Microscopy, optical tweezers, quantitative phase imaging, dynamic light scattering, ektacytometry, filtration and microfluidic devices have been used to study the behaviour of individual RBCs and populations of RBCs(Kim *et al*., 2012). These different techniques show different sensitivities to contributions from RBC geometry, cytoplasmic viscosity and plasma membrane viscoelasticity, making it difficult to correlate *in vitro* measures of biomechanical properties with clinical pathologies(Ataga *et al*., 2011, Huisjes *et al*., 2018, Nader *et al*., 2019).

In this work, we examined the individual contributions of cellular rigidity and geometry on the ability of RBCs to conform into microchannels. We made use of a modified version of the HEMA microfluidic device(Gifford *et al*., 2003) in which RBCs are flowed into wedge-shaped microchannels. Conforming RBCs into the microchannels allows estimation of the surface area, the volume and the MCD, *i*.*e*. the diameter of the smallest equivalent cylindrical tube through which an RBC could pass. Small variations in channel geometry were observed between different HEMA chips and, where possible, we examined different samples in the same chip by differentially labelling one population or washing and reusing the chip for different samples on the same day.

We altered the SA:V ratio of RBCs by modifying the buffer osmolarity. In agreement with previous reports(Reinhart *et al*., 1985, Gifford *et al*., 2003, Safeukui *et al*., 2012), we found that the ability of RBCs to travel into microchannels is highly sensitive to the SA:V ratio. Our experiments revealed a roughly inverse relationship between MCD and SA:V ratio. Importantly, in the SA:V range close to physiological levels, *i*.*e*. 1.38 to 1.58, the 15% increase in MCD was readily detected by HEMA. By contrast, we found that ektacytometry, a method that is often used to measure RBC defects, was insensitive to changes in the SA:V ratio over this range.

Somewhat surprisingly, substantial increases in RBC stiffness did not affect the ability of RBCs to progress into the channels. For this work, we used glutaraldehyde, a bifunctional linker that cross-links membrane proteins, thereby decreasing the shear modulus, as well as polymerising haemoglobin, thus increasing cytoplasmic viscosity. Treatment with 0.004% glutaraldehyde largely abrogated the ability of RBCs to elongate in fluid flow. Such changes in the rheological properties of the RBCs would be expected to markedly increase blood viscosity at high shear stress, which would adversely affect the rheology of bulk flow(Mohandas *et al*., 2008, Pivkin *et al*., 2016). By contrast, these highly rigidified cells migrated to a similar position in the microchannels as untreated RBCs. These data suggest that cellular rigidity plays a less important role in microcirculatory blood flow than generally appreciated.

To examine the consequences of our findings for different pathological conditions, we examined the biomechanical properties of RBCs infected with two different species of malaria parasite. We found that RBCs infected with *P. falciparum* in the trophozoite stage of infection are highly rigidified, exhibiting no elongation in fluid flow, even at the highest applied shear stress, similar to the behaviour of RBCs treated with the highest glutaraldehyde concentration. This is likely due to the greatly increased shear modulus of the host RBC membrane(Suresh *et al*., 2005, Park *et al*., 2008), as well as the presence of a rigid intracellular parasite, which occupies about 60% of the host cell cytoplasm(Hanssen *et al*., 2012). In our work, we found that *P. falciparum* trophozoite-infected RBCs maintained the same SA:V ratio as uninfected RBCs. Some other studies have reported that *P. falciparum*-infected RBCs exhibit a decreased SA:V ratio(Herricks *et al*., 2009, Safeukui *et al*., 2012). This discrepancy may arise from differences in the methods used to assess SA:V ratio, or from the fact that previous studies examined RBCs infected with more mature schizont stage parasites, which undergo permeability changes leading up to parasite egress(Hale *et al*., 2017). Our work suggests that the rigidification of trophozoite-stage infected RBCs is not, in and of itself, sufficient to hinder passage through microcapillaries. Nonetheless, rigidification of the RBC membrane will enhance adhesion to microvessel walls by distributing the tensional forces imposed on individual adhesin molecules through to the membrane skeletal network(Zhang *et al*., 2015). Adherent *P. falciparum*-infected RBCs may narrow the blood vessels, leading to increased microvascular obstruction and downstream clinical complications(Hanson *et al*., 2015).

Our results have important implications for *P. knowlesi* infections. In Malaysia, *P. knowlesi* now accounts for >90% of all malaria cases(Cooper *et al*., 2020). Severe *P. knowlesi* malaria is associated with haemolysis-induced endothelial activation leading to acute kidney injury(Barber *et al*., 2018a, Grigg *et al*., 2018). We found that *P. knowlesi*-infected RBCs exhibit a large increase in volume and a more moderate increase in surface area, resulting in an overall decrease in SA:V ratio and a consequent increase in MCD. We anticipate that the increased MCD may underpin the ability of *P. knowlesi*-infected cells to sequester within the vasculature despite the absence of surface-exposed adhesins(Knisely *et al*., 1964, Menezes *et al*., 2012). The consequent evasion of the splenic clearance mechanisms may enable increased parasite density. The increased volume may also underpin the finding that *P. knowlesi*-infected RBCs are hyper-sensitive to hypotonic lysis(Liu *et al*., 2019). Chronic intravascular rupture of swollen *P. knowlesi*-infected RBCs may lead to chronic release of haem and iron(Van Avondt *et al*., 2019), which may in turn lead to the acute kidney injury that characterises severe knowlesi malaria.

We also examined the geometry and biomechanical properties of early stage reticulocytes. When first released from the bone marrow, reticulocytes contain RNA and have remnant surface receptors, particularly the transferrin receptor, CD71, which can be labelled with antibodies. As the reticulocyte matures, CD71 is lost via a process of shedding and endocytosis. In the final stages of maturation the RNA is degraded(Malleret *et al*., 2013, Mankelow *et al*., 2015). We used CD71 labelling of density gradient-purified reticulocytes to identify early stage reticulocytes and found that they have 30% more surface area than mature RBCs. This difference is larger than reported in previous studies that detected reticulocytes based on RNA staining(Waugh *et al*., 1997, Gifford *et al*., 2006). Our data confirm that a loss of surface area accompanies the loss of surface markers. In agreement with previous studies we found that reticulocytes exhibit a decreased ability to elongate in fluid flow. Nonetheless, even the early stage (CD71+) reticulocytes exhibit a very similar SA:V ratio to mature RBCs, which likely underpins their ability to passage through the microcirculation without disturbing blood flow.

We generated a nonlinear 3D FE model to further interrogate the biomechanical basis for our surprising finding that cell stiffness makes very little contribution to the ability of RBCs to traverse small channels. These simulations are consistent with our experimental observation. The RBC membrane skeleton imparts extremely nonlinear mechanical behaviour(Mills *et al*., 2004), which means that the RBC can be stretched sufficiently to conform to the microchannel even when the cellular rigidity increases. At very high shear modulus values, we observed the formation of creases in the membrane that prevent the RBC from travelling into smaller equivalent diameters, as strain and stress in the membrane are significantly increased in the vicinity of the creases. Importantly, our simulations are consistent with the experimental finding that a moderate decrease in the SA:V ratio, *i*.*e*. moderately increased RBC sphericity, imposes a high deformation on the RBC and markedly affects its ability to traverse small channels. We note, as a limitation of our study, that we did not consider time-dependent parameters, such as the rate of progress of RBCs in the microchannels, prior to being trapped.

We were able to use our FE model to predict cut-off values for shear modulus and SA:V ratio that would prevent RBCs passing through small channels with diameters comparable to cerebral cortex capillaries. These values are difficult to obtain experimentally. Our analysis reveals that stiffening the RBCs has relatively little effect on the minimum applied pressure needed to force RBCs to passage through capillaries, even of 3 μm diameter. Thus, even highly rigidified RBCs would be expected to passage through the microcirculation, so long as the SA:V ratio is kept constant. By contrast, reducing the SA:V ratio to 1.2 increases the minimum applied pressure by ∼15-fold. We note that while our work provides insights into the determinants of passage of modified RBCs through small capillaries, they do not address the biomechanical parameters that permit passage through splenic endothelial slits, which can be as narrow as 1 – 2 μm. Further work is needed to measure and model the individual effects of cellular rigidity and SA:V ratio of this important biomechanical surveillance mechanism.

In summary, we have employed a modified HEMA microfluidic device that enables accurate determination of volume, surface area and the capillary diameter through which an RBC can passage. We found that RBC rigidity makes very little contribution to RBC behaviour in microchannels due to the nonlinear mechanical behaviour of the RBC membrane skeleton. Our work has particular implications for diseases such as hereditary spherocytosis and hereditary elliptocytosis(Warkentin *et al*., 1990, Da Costa *et al*., 2016) that exhibit a decreased SA:V ratio. Following splenectomy – a treatment that is commonly used to reduce anaemia – circulation of RBCs with decreased SA:V ratio may impair blood flow through small capillaries, contributing to an enhanced risk of thrombosis(Cappellini *et al*., 2000, Schilling *et al*., 2008, Byrnes *et al*., 2017, Iolascon *et al*., 2017). Application of techniques that are highly sensitive to changes is SA:V ratio may provide an important indicator of the pathological consequences of changes in RBC geometry.

## Experimental procedures

### Glutaraldehyde treatment

Glutaraldehyde (Sigma, EM grade) solutions in PBS were prepared at the required concentration. O+ human RBCs (Red Cross Blood Band, Australia) were gently dispersed at 2% haematocrit and incubated at room temperature (RT) for 1 h. The RBCs were pelleted at 300 g for 2 min, resuspended in PBS and washed three times. The treated cells were mixed at a 1:1 ratio with untreated cells from the same RBC aliquot before introduction to the microfluidic device. Treated cells were identified by their autofluorescence in long-exposure images collected with 632 nm excitation/ 676 (34) nm emission filter sets.

### Altering buffer osmolarity

The buffer osmolarity was tuned over the range 150-520 mOsm/kg by adjusting the NaCl content while keeping the Na_2_HPO_4_, KH_2_PO_4_ and KCl concentrations constant at 10, 1.8 and 2.7 mM respectively. The pH was adjusted to 7.35 before measurement of osmolarity using an Advanced Instruments 3320 freezing-point osmometer. RBCs were washed once in PBS before being dispersed in osmolarity-modified PBS at 2% haematocrit. RBC suspensions were then further diluted with modified PBS before introduction to the microfluidic device.

### Culture of *P. falciparum* and *P. knowlesi*

*P. falciparum* (3D7 strain) and *P. knowlesi* (A1 strain) were cultured as described previously(Liu *et al*., 2019). Briefly, parasites were maintained in O+ human RBCs (5% haematocrit) in RPMI-based complete medium containing pooled human serum (5%), AlbuMAX™ II (5%), 200 μM hypoxanthine, 10 mM D-glucose (Sigma) and 20 μg/ml gentamicin (Sigma). Parasitemia was routinely maintained below 5%. Synchronized trophozoite-stage infected RBCs (32-36 h post-invasion *P. falciparum*; 22 – 26 h post-invasion *P. knowlesi*) were enriched from culture by magnetic separation(Liu *et al*., 2019).

### Reticulocyte Purification

500 ml of whole blood (Australian Red Cross) or 80 ml cord blood (BMDI Cord Blood Bank) was passed through an RC High Efficiency leucocyte removal filter (Haemonetics Australia) and washed three times with PBS to remove serum components. Donated blood was approved for use by the Walter and Eliza Hall Institute Human Research Ethics Committee (HREC 14/09). RBCs at approximately 50% haematocrit were layered onto a 70% (v/v) isotonic Percoll cushion (GE Healthcare) and centrifuged for 25 min at 2,000 x g. The thin band formed at the Percoll interface was collected as the reticulocyte fraction, while the pellet was collected as the mature cell fraction. The percentage of reticulocytes in the reticulocyte fraction was measured by flow cytometry. To stain reticulocytes, red blood cells in a concentration of 1 × 10^7^ cells/mL were incubated for half an hour with 100 μl thiazole orange (TO) (Retic-Count Reagent; BD Biosciences) and Alexa-647 conjugated mouse mAb against transferrin receptor (CD71, mouse mAb MEM75, Abcam ab187777). The red blood cells were washed with PBS then resuspended in 200 μl PBS and analysed on the FACSCalibur flow cytometer (BD Biosciences). 50,000 events were recorded and results were analysed using FlowJo software (Three Star). Cells were stored in RPMI wash buffer (Gibco) at 4°C until needed. Reticulocytes were identified by labelling with antisera for CD71 (mouse mAb 13E4, Abcam ab38171) and anti-mouse Alexa-488 secondary. Pelleted (mature) RBCs were tagged by labelling with antisera against complement receptor-1 (CD35, mouse mAb543), and anti-mouse Alexa 647 secondary. Labelling was performed at 1:500 in PBS solution with 3% bovine serum albumin (BSA) for 1 h at RT followed by at least 3 washing steps in PBS. Mature and immature cell types were mixed at a ratio of 1:3 following labelling before introduction to the HEMA device. Unlabelled cells were considered to be CD71-reticulocytes, however this fraction may also include unlabelled (CD35-) mature cells.

### Ektacytometry

An aliquot of packed RBCs (10 μL) was dispersed into 500 μL of 6% 360 kDa polyvinylpyrrolidone in PBS. The elongation index (EI) was measured using a RheoScan ektacytometer(Shin *et al*., 2005, Dearnley *et al*., 2016), at stress stresses of 0–20 Pa. Each measurement was performed in triplicate and the averaged data are presented. Magnet-purified packed RBCs (parasitemia >97%) were used for the analysis of infected RBCs.

### Fabrication of HEMA microfluidic chips

Our microchannel arrays are based on a modified original design(Gifford *et al*., 2003) and are fabricated using standard soft lithography procedure(Madou, 2002, Lake *et al*., 2015). Briefly, polydimethyl-siloxane (PDMS) elastomer is applied to a pre-fabricated master mould (Melbourne Centre for Nanofabrication) and cured in an oven. Holes punched into the PDMS before glass coverslip attachment allow for connection of tubing to a fluid reservoir and pressure pump. The microchannels were 125.2 ± 0.06 μm long and 3.23 ± 0.05 μm deep as measured via SEM. The width of wedge-shaped microchannels was 4.98 ± 0.07 μm at the entrance and 1.44 ± 0.04 μm at the exit. Small variations in geometry were observed between different HEMA chips.

### SEM Imaging

PDMS containing the HEMA pattern was cut to size and into cross-sections with a razor blade before mounting on conducting carbon tape. PDMS sections were sputter coated from a gold target to a thickness of ∼0.5 nm before measurement. Reticulocytes and mature RBCs were pre-fixed in 0.005% glutaraldehyde in PBS for 20 min at RT before full fixation in 2.5% glutaraldehyde in PBS for 1.5 h at RT. Agarose-embedded cells were stained in ferrocyanide-reduced osmium tetroxide in 0.15 M cacodylate buffer for 1 h on ice. After washing, the preparation was incubated with freshly prepared 1% thiocarbonhydrazide solution in H_2_O for 20 min at RT, stained with 2% osmium tetroxide in H_2_O for 30 min at RT, stained with 1% aqueous uranyl acetate overnight followed by Walton’s lead aspartate for 30 min at 60°C. The preparation was dehydrated in a graded series of ethanol-H_2_O, followed by progressive infiltration with EPON resin. After polymerization, the resin block was trimmed, mounted on a microtome stub using silver glue on an ultramicrotome (UC7, Leica). The resin block face was polished again with a diamond knife on the ultramicrotome after 10 nm gold coating. Serial images were collected using a serial block face-scanning electron microscope (SBF-SEM), equipped with an in-chamber diamond knife (Teneo VolumeScope, FEI Company), using the back-scattered electron signal at 3 kV, under low vacuum conditions. The section thickness was set to 50 nm and the pixel size of each image in the stack was 5 nm. Serial sections were contrasted and aligned using IMOD software (Boulder Laboratory for 3D Electron Microscopy of Cells). Raw images were binned to 20 nm and smoothed with a Gaussian function before segmentation using contours and thresholding.

### HEMA experimental setup

The HEMA chip was purged (using a syringe pump) with 80% ethanol at a flow rate of 2 μL/min for 30 min, then flushed with PBS or complete culture media for 45 min. Diluted washed blood (at 0.2-0.5% haematocrit) was driven into the microchannels at a flow pressure of 180-250 mbar using an Elveflow AF1 pressure pump. Experiments were completed within 1 h, as infected cells trapped in the device were prone to lysis. Due to a slight variability in dimensions from device to device, for all experiments except the glutaraldehyde study, the same microchip was used for several samples. The microfluidic channels were backflushed with PBS for 20 mins at a flow rate of 3 μL/min before flushing with the new buffer for 30 min between each sample. Microfluidic chips were mounted in a Deltavision Elite (GE Healthcare) inverted microscope and imaged using a 40x air objective at 37°C for malaria parasite-infected cells and at RT for other experiments. Images were collected in brightfield along with the blue fluorescence channel (390/18 nm excitation and 435/48 nm collection filter sets) to facilitate automated analysis using the shadow of RBC haemoglobin in the fluorescence images.

Acquired images were analysed in ImageJ (NIH Image/ImageJ) and cells were delineated automatically using a macro plugin developed in-house. Volume, surface area, SA:V ratio and MCD were calculated by importing the measurements into R software.

### MCD calculation

MCD is calculated as the cross-sectional area (Area_m_) of an arrested RBC at its midpoint (Eq. 1).

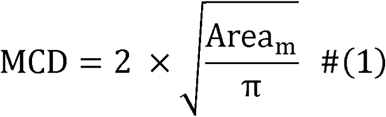

### 3D FE model

The RBC membrane was discretised using four-node linear shell elements, and the third order Yeoh hyper-elastic model was used to represent the stress-strain relationships in the membrane. For all computations, a typical bending modulus of 2×10^−19^ Nm was assumed(Mills *et al*., 2004, Yoon *et al*., 2016). Cytosol, an incompressible fluid inside the membrane, keeps the cell volume constant under the deformation. Thus, we regarded the cytosol as a volume constant constraint inside the cell membrane and ran all the simulations using the quasi-static analysis. Blood vessel walls are reported to be two to three orders of magnitude stiffer than RBCs(Xie *et al*., 1995, Lu *et al*., 2003). Therefore, we ignored deformation of the capillary wall and regarded the capillaries as rigid parts in the FE model.

### Statistics and reproducibility

Statistical analysis employed the Welch two samples t-test (R software). Each of the presented dataset are representative of or averages of data obtained during at least two different experimental sessions, using different batches of blood, as detailed in the text.

### Data availability

The authors main data supporting the findings in this study are provided in this article (and the supplementary information). Individual data points for the different HEMA analysis sets are available upon request.

## Acknowledgements

We thank the Australian Red Cross Blood Service for whole blood. Cord blood units were supplied by the BMDI Cord Blood Bank at the Murdoch Childrens Research Institute and The Royal Children’s Hospital. Establishment and running of the BMDI Cord Blood Bank is made possible through generous support by Federal and state governments, the Murdoch Childrens Research Institute, The Royal Children’s Hospital Foundation and the Fight Cancer Foundation. We thank Sebastien Menant, the Walter and Eliza Hall Institute for help with preparation of reticulocytes. Light microscopy was performed at the Biological Optical Microscopy Platform and electron microscopy at the Bio21 Institute Advanced Microscopy Facility, The University of Melbourne (www.microscopy.unimelb.edu.au). This work was performed in part at the Melbourne Centre for Nanofabrication (MCN) in the Victorian Node of the Australian National Fabrication Facility (ANFF).

## Author contributions

L.T., V.R., A.N, A.J.B, P.V.S.L. and M.W.D. designed the study; A.N, A.J.B. and B.L. performed the experiments; A,N, A.J.B, P.V.S.L. and M.W.D. analysed the data. O.C., S.T, O.L., D.A., J-L. C., and W-H.T. characterized and provided the samples; L.T., V.R., A.N, A.J.B, P.V.S.L. and M.W.D wrote the paper. All authors participated in discussion and paper editing.

## Additional information

Supplementary information accompanies this paper.

## References

Aingaran, M., Zhang, R., Law, S.K., Peng, Z., Undisz, A., Meyer, E., et al. (2012). Host cell deformability is linked to transmission in the human malaria parasite *Plasmodium falciparum*. Cell Microbiol 14, 983–993.

Allison, A.C. (1960). Turnovers of erythrocytes and plasma proteins in mammals. Nature 188, 37–40.

An, X. and Mohandas, N. (2008). Disorders of red cell membrane. Br J Haematol 141, 367–375.

Ataga, K.I., Reid, M., Ballas, S.K., Yasin, Z., Bigelow, C., James, L.S., et al. (2011). Improvements in haemolysis and indicators of erythrocyte survival do not correlate with acute vaso-occlusive crises in patients with sickle cell disease: a phase III randomized, placebo-controlled, double-blind study of the Gardos channel blocker senicapoc (ICA-17043). Br J Haematol 153, 92–104.

Barber, B.E., Grigg, M.J., Piera, K.A., William, T., Cooper, D.J., Plewes, K., et al. (2018a). Intravascular haemolysis in severe *Plasmodium knowlesi* malaria: association with endothelial activation, microvascular dysfunction, and acute kidney injury. Emerg Microbes Infect 7, 106.

Barber, B.E., Russell, B., Grigg, M.J., Zhang, R., William, T., Amir, A., et al. (2018b). Reduced red blood cell deformability in *Plasmodium knowlesi* malaria. Blood Adv 2, 433–443.

Barber, B.E., William, T., Grigg, M.J., Menon, J., Auburn, S., Marfurt, J., et al. (2013). A prospective comparative study of *knowlesi, falciparum*, and *vivax* malaria in Sabah, Malaysia: high proportion with severe disease from Plasmodium *knowlesi* and Plasmodium *vivax* but no mortality with early referral and artesunate therapy. Clin Infect Dis 56, 383–397.

Beare, N.A., Harding, S.P., Taylor, T.E., Lewallen, S. and Molyneux, M.E. (2009). Perfusion abnormalities in children with cerebral malaria and malarial retinopathy. J Infect Dis 199, 263–271.

Byrnes, J.R. and Wolberg, A.S. (2017). Red blood cells in thrombosis. Blood 130, 1795–1799.

Canham, P.B. and Burton, A.C. (1968). Distribution of size and shape in populations of normal human red cells. Circ Res 22, 405–422.

Cappellini, M.D., Robbiolo, L., Bottasso, B.M., Coppola, R., Fiorelli, G. and Mannucci, A.P. (2000). Venous thromboembolism and hypercoagulability in splenectomized patients with thalassaemia intermedia. Br J Haematol 111, 467–473.

Chang, C.Y., Pui, W.C., Kadir, K.A. and Singh, B. (2018). Spontaneous splenic rupture in *Plasmodium knowlesi* malaria. Malar J 17, 448.

Clark, M., Mohandas, N. and Shohet, S. (1983). Osmotic gradient ektacytometry: comprehensive characterization of red cell volume and surface maintenance. Blood 61, 899–910.

Cooper, D.J., Rajahram, G.S., William, T., Jelip, J., Mohammad, R., Benedict, J., et al. (2020). *Plasmodium knowlesi* malaria in Sabah, Malaysia, 2015-2017: Ongoing increase in incidence despite near-elimination of the human-only *Plasmodium* species. Clin Infect Dis 70, 361–367.

Cox-Singh, J. and Singh, B. (2008). Knowlesi malaria: newly emergent and of public health importance? Trends Parasitol 24, 406–410.

Da Costa, L., Suner, L., Galimand, J., Bonnel, A., Pascreau, T., Couque, N., et al. (2016). Diagnostic tool for red blood cell membrane disorders: Assessment of a new generation ektacytometer. Blood Cells Mol Dis 56, 9–22.

de Koning-Ward, T.F., Dixon, M.W., Tilley, L. and Gilson, P.R. (2016). Plasmodium species: master renovators of their host cells. Nat Rev Microbiol 14, 494–507.

Dearnley, M., Chu, T., Zhang, Y., Looker, O., Huang, C., Klonis, N., et al. (2016). Reversible host cell remodeling underpins deformability changes in malaria parasite sexual blood stages. Proc Natl Acad Sci U S A 113, 4800–4805.

Diez-Silva, M., Dao, M., Han, J., Lim, C.T. and Suresh, S. (2010). Shape and biomechanical characteristics of human red blood cells in health and disease. MRS Bull 35, 382–388.

Dondorp, A.M., Kager, P.A., Vreeken, J. and White, N.J. (2000). Abnormal blood flow and red blood cell deformability in severe malaria. Parasitology Today 16, 228–232.

Evans, E. and Fung, Y.-C. (1972). Improved measurements of the erythrocyte geometry. Microvascular Research 4, 335–347.

Evans, E., Mohandas, N. and Leung, A. (1984). Static and dynamic rigidities of normal and sickle erythrocytes. Major influence of cell hemoglobin concentration. The Journal of clinical investigation 73, 477–488.

Forsyth, A.M., Wan, J., Ristenpart, W.D. and Stone, H.A. (2010). The dynamic behavior of chemically “stiffened” red blood cells in microchannel flows. Microvasc Res 80, 37–43.

Gifford, S.C., Derganc, J., Shevkoplyas, S.S., Yoshida, T. and Bitensky, M.W. (2006). A detailed study of time-dependent changes in human red blood cells: from reticulocyte maturation to erythrocyte senescence. Br J Haematol 135, 395–404.

Gifford, S.C., Frank, M.G., Derganc, J., Gabel, C., Austin, R.H., Yoshida, T. and Bitensky, M.W. (2003). Parallel microchannel-based measurements of individual erythrocyte areas and volumes. Biophys J 84, 623–633.

Glenister, F.K., Coppel, R.L., Cowman, A.F., Mohandas, N. and Cooke, B.M. (2002). Contribution of parasite proteins to altered mechanical properties of malaria-infected red blood cells. Blood 99, 1060–1063.

Grigg, M.J., William, T., Barber, B.E., Rajahram, G.S., Menon, J., Schimann, E., et al. (2018). Age-related clinical spectrum of *Plasmodium knowlesi* malaria and predictors of severity. Clin Infect Dis 67, 350–359.

Hale, V.L., Watermeyer, J.M., Hackett, F., Vizcay-Barrena, G., van Ooij, C., Thomas, J.A., et al. (2017). Parasitophorous vacuole poration precedes its rupture and rapid host erythrocyte cytoskeleton collapse in *Plasmodium falciparum* egress. Proc Natl Acad Sci U S A 114, 3439–3444.

Hanson, J., Lee, S.J., Hossain, M.A., Anstey, N.M., Charunwatthana, P., Maude, R.J., et al. (2015). Microvascular obstruction and endothelial activation are independently associated with the clinical manifestations of severe falciparum malaria in adults: an observational study. BMC Med 13, 122.

Hanssen, E., Knoechel, C., Dearnley, M., Dixon, M.W., Le Gros, M., Larabell, C. and Tilley, L. (2012). Soft X-ray microscopy analysis of cell volume and hemoglobin content in erythrocytes infected with asexual and sexual stages of *Plasmodium falciparum*. J Struct Biol 177, 224–232.

Herricks, T., Antia, M. and Rathod, P.K. (2009). Deformability limits of *Plasmodium falciparum*-infected red blood cells. Cell Microbiol 11, 1340–1353.

Hochmuth, R.M. and Waugh, R.E. (1987). Erythrocyte Membrane Elasticity and Viscosity. Annual Review of Physiology 49, 209–219.

Huisjes, R., Bogdanova, A., van Solinge, W.W., Schiffelers, R.M., Kaestner, L. and van Wijk, R. (2018). Squeezing for life - Properties of red blood cell deformability. Front Physiol 9, 656.

Iolascon, A., Andolfo, I., Barcellini, W., Corcione, F., Garçon, L., De Franceschi, L., et al. (2017). Recommendations regarding splenectomy in hereditary hemolytic anemias. Haematologica 102, 1304–1313.

Kim, Y., Kim, K. and Park, Y. (2012) Measurement techniques for red blood cell deformability: Recent advances. In Blood Cell - An Overview of Studies in Hematology, T.E. Moschandreou (ed.).

Knisely, M.H., Stratman-Thomas, W.K., Eliot, T.S. and Bloch, E.H. (1964). Knowlesi malaria in monkeys II: A first step in the separation of the mechanical pathologic circulatory factors of one sludge disease from possible specific toxic factors of that disease. Angiology 15, 411–416.

Kono, M., Kondo, T., Takagi, Y., Wada, A. and Fujimoto, K. (2009). Morphological definition of CD71 positive reticulocytes by various staining techniques and electron microscopy compared to reticulocytes detected by an automated hematology analyzer. Clin Chim Acta 404, 105–110.

Kuhn, V., Diederich, L., Keller, T.C.S.t., Kramer, C.M., Luckstadt, W., Panknin, C., et al. (2017). Red blood cell function and dysfunction: Redox regulation, nitric oxide metabolism, anemia. Antioxid Redox Signal 26, 718–742.

Lake, M., Narciso, C., Cowdrick, K., Storey, T., Zhang, S., Zartman, J. and Hoelzle, D.J.P.E. (2015). Microfluidic device design, fabrication, and testing protocols. 10.

Lauwers, F., Cassot, F., Lauwers-Cances, V., Puwanarajah, P. and Duvernoy, H. (2008). Morphometry of the human cerebral cortex microcirculation: general characteristics and space-related profiles. Neuroimage 39, 936–948.

Lelliott, P.M., Huang, H.M., Dixon, M.W., Namvar, A., Blanch, A.J., Rajagopal, V., et al. (2017). Erythrocyte beta spectrin can be genetically targeted to protect mice from malaria. Blood Adv 1, 2624–2636.

Li, H., Yang, J., Chu, T.T., Naidu, R., Lu, L., Chandramohanadas, R., et al. (2018). Cytoskeleton remodeling induces membrane stiffness and stability changes of maturing reticulocytes. Biophys J 114, 2014–2023.

Linninger, A.A., Gould, I.G., Marrinan, T., Hsu, C.Y., Chojecki, M. and Alaraj, A. (2013). Cerebral microcirculation and oxygen tension in the human secondary cortex. Annals of biomedical engineering 41, 2264–2284.

Liu, B., Blanch, A.J., Namvar, A., Carmo, O., Tiash, S., Andrew, D., et al. (2019). Multimodal analysis of *Plasmodium knowlesi*-infected erythrocytes reveals large invaginations, swelling of the host cell, and rheological defects. Cell Microbiol, e13005.

Liu, J., Guo, X., Mohandas, N., Chasis, J.A. and An, X. (2010). Membrane remodeling during reticulocyte maturation. Blood 115, 2021–2027.

Lu, X., Yang, J., Zhao, J.B., Gregersen, H. and Kassab, G.S. (2003). Shear modulus of porcine coronary artery: contributions of media and adventitia. American Journal of Physiology-Heart and Circulatory Physiology 285, H1966–H1975.

Madou, M.J. (2002) Fundamentals of microfabrication: the science of miniaturization, CRC press.

Malleret, B., Xu, F., Mohandas, N., Suwanarusk, R., Chu, C., Leite, J.A., et al. (2013). Significant biochemical, biophysical and metabolic diversity in circulating human cord blood reticulocytes. PLoS One 8, e76062.

Mankelow, T.J., Griffiths, R.E., Trompeter, S., Flatt, J.F., Cogan, N.M., Massey, E.J. and Anstee, D.J. (2015). Autophagic vesicles on mature human reticulocytes explain phosphatidylserine-positive red cells in sickle cell disease. Blood 126, 1831–1834.

Mankelow, T.J., Satchwell, T.J. and Burton, N.M. (2012). Refined views of multi-protein complexes in the erythrocyte membrane. Blood Cells Mol Dis 49, 1–10.

Marin-Padilla, M. (2012). The human brain intracerebral microvascular system: development and structure. Front Neuroanat 6, 38.

Menezes, R.G., Pant, S., Kharoshah, M.A., Senthilkumaran, S., Arun, M., Nagesh, K.R., et al. (2012). Autopsy discoveries of death from malaria. Leg Med (Tokyo) 14, 111–115.

Mills, J.P., Qie, L., Dao, M., Lim, C.T. and Suresh, S. (2004). Nonlinear elastic and viscoelastic deformation of the human red blood cell with optical tweezers. Mech Chem Biosyst 1, 169–180.

Mohandas, N. and Gallagher, P.G. (2008). Red cell membrane: past, present, and future. Blood 112, 3939–3948.

Nader, E., Skinner, S., Romana, M., Fort, R., Lemonne, N., Guillot, N., et al. (2019). Blood rheology: Key parameters, impact on blood flow, role in sickle cell disease and effects of exercise. Front Physiol 10, 1329.

Park, Y., Diez-Silva, M., Popescu, G., Lykotrafitis, G., Choi, W., Feld, M.S. and Suresh, S. (2008). Refractive index maps and membrane dynamics of human red blood cells parasitized by *Plasmodium falciparum*. Proc Natl Acad Sci U S A 105, 13730–13735.

Pivkin, I.V., Peng, Z., Karniadakis, G.E., Buffet, P.A., Dao, M. and Suresh, S. (2016). Biomechanics of red blood cells in human spleen and consequences for physiology and disease. Proceedings of the National Academy of Sciences 113, 7804–7809.

Reinhart, W.H. and Chien, S. (1985). Roles of cell geometry and cellular viscosity in red cell passage through narrow pores. Am J Physiol 248, C473–479.

Renia, L., Howland, S.W., Claser, C., Charlotte Gruner, A., Suwanarusk, R., Hui Teo, T., et al. (2012). Cerebral malaria: mysteries at the blood-brain barrier. Virulence 3, 193–201.

Renoux, C., Faivre, M., Bessaa, A., Da Costa, L., Joly, P., Gauthier, A. and Connes, P. (2019). Impact of surface-area-to-volume ratio, internal viscosity and membrane viscoelasticity on red blood cell deformability measured in isotonic condition. Sci Rep 9, 6771.

Safeukui, I., Buffet, P.A., Deplaine, G., Perrot, S., Brousse, V., Ndour, A., et al. (2012). Quantitative assessment of sensing and sequestration of spherocytic erythrocytes by the human spleen. Blood 120, 424–430.

Safeukui, I., Buffet, P.A., Deplaine, G., Perrot, S., Brousse, V., Sauvanet, A., et al. (2018). Sensing of red blood cells with decreased membrane deformability by the human spleen. Blood Adv 2, 2581–2587.

Schilling, R.F., Gangnon, R.E. and Traver, M.I. (2008). Delayed adverse vascular events after splenectomy in hereditary spherocytosis. Journal of thrombosis and haemostasis : JTH 6, 1289–1295.

Shin, S., Ku, Y., Park, M.-S. and Suh, J.-S. (2005). Slit-flow ektacytometry: Laser diffraction in a slit rheometer. Cytometry Part B: Clinical Cytometry 65B, 6–13.

Singh, B. and Daneshvar, C. (2013). Human infections and detection of *Plasmodium knowlesi*. Clin Microbiol Rev 26, 165–184.

Suresh, S., Spatz, J., Mills, J.P., Micoulet, A., Dao, M., Lim, C.T., et al. (2005). Connections between single-cell biomechanics and human disease states: gastrointestinal cancer and malaria. Acta Biomaterialia 1, 15–30.

Suzuki, Y., Tateishi, N., Soutani, M. and Maeda, N. (1996). Flow behavior of erythrocytes in microvessels and glass capillaries: effects of erythrocyte deformation and erythrocyte aggregation. Int J Microcirc Clin Exp 16, 187–194.

Tomaiuolo, G. (2014). Biomechanical properties of red blood cells in health and disease towards microfluidics. Biomicrofluidics 8, 051501.

Van Avondt, K., Nur, E. and Zeerleder, S. (2019). Mechanisms of haemolysis-induced kidney injury. Nat Rev Nephrol 15, 671–692.

Wahlgren, M., Goel, S. and Akhouri, R.R. (2017). Variant surface antigens of *Plasmodium falciparum* and their roles in severe malaria. Nat Rev Microbiol 15, 479–491.

Warkentin, T.E., Barr, R.D., Ali, M.A. and Mohandas, N. (1990). Recurrent acute splenic sequestration crisis due to interacting genetic defects: hemoglobin SC disease and hereditary spherocytosis. Blood 75, 266–270.

Warrell, D.A., White, N.J., Veall, N., Looareesuwan, S., Chanthavanich, P., Phillips, R.E., et al. (1988). Cerebral anaerobic glycolysis and reduced cerebral oxygen transport in human cerebral malaria. Lancet 2, 534–538.

Waugh, R.E., McKenney, J.B., Bauserman, R.G., Brooks, D.M., Valeri, C.R. and Snyder, L.M. (1997). Surface area and volume changes during maturation of reticulocytes in the circulation of the baboon. J Lab Clin Med 129, 527–535.

Xie, J., Zhou, J. and Fung, Y.C. (1995). Bending of Blood Vessel Wall: Stress-Strain Laws of the Intima-Media and Adventitial Layers. Journal of Biomechanical Engineering 117, 136–145.

Yeoh, O.H. (1990). Characterization of elastic properties of carbon-black-filled rubber vulcanizates. Rubber Chemistry and Technology 63, 792–805.

Yoon, D. and You, D. (2016). Continuum modeling of deformation and aggregation of red blood cells. Journal of biomechanics 49, 2267–2279.

Zhang, Y., Huang, C., Kim, S., Golkaram, M., Dixon, M.W., Tilley, L., et al. (2015). Multiple stiffening effects of nanoscale knobs on human red blood cells infected with *Plasmodium falciparum* malaria parasite. Proc Natl Acad Sci U S A 112, 6068–6073.

