## Supplementary Information for "Surface area-to-volume ratio, not cellular rigidity, determines red blood cell traversal through small capillaries"

**Cellular Microbiology**

**Supplementary Figure 1.** **Scanning electron microscopic (SEM) images of the Human Erythrocyte Microchannel Analyser (HEMA) polydimethylsiloxane (PDMS) chip.** The microchannels are 125.2 $\pm$0.06 μm in length and 3.23 $\pm$0.05 μm deep. The width of the wedge-shaped microchannels is 4.98 $\pm$0.07 μm at the entrance and 1.44$\pm$0.04 μm at the exit.

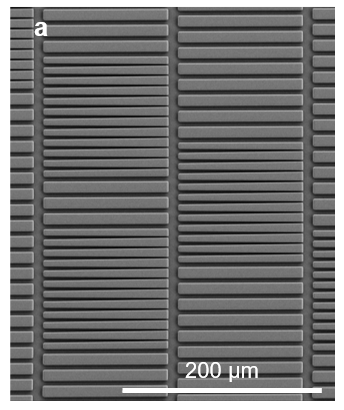

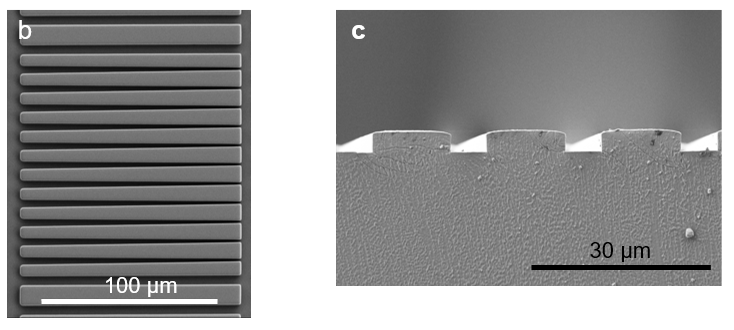

**Supplementary Figure 2. SA:V ratio and MCD for unfixed and fixed red blood cells (RBCs).**

Comparison of geometric parameters for unfixed RBCs (white) and fixed RBCs (grey). (a) Values of SA:V ratio for the experiment depicted in Fig 1d,e. (b,c) Values of SA:V ratio and MCD for an independent experiment. Fixed RBCs at each glutaraldehyde concentration were mixed with unfixed RBCs before analysis on an individual HEMA chip. Small variations in channel geometry were observed between different HEMA chips. Comparisons are made between RBCs in the same chip, *i.e.* grey and white bars. Mean values and first and third quartiles are depicted.

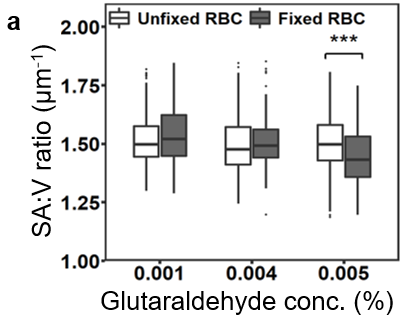

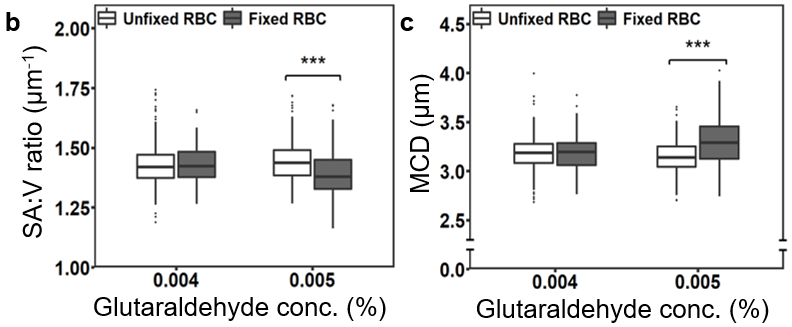

**Supplementary Figure 3. No creases are present in the membrane of trapped normal RBCs.**

Healthy RBCs were labelled with PKH67 dye following the manufacturer’s protocol before introduction into the HEMA chip. A Z stack of a trapped RBC was acquired using the green channel of a DeltaVision Elite fluorescence microscope using a 100x oil objective. No creases or folds in the membrane were observed in 55 Z-planes with the nominal spacing of $0.2 \mu m$.

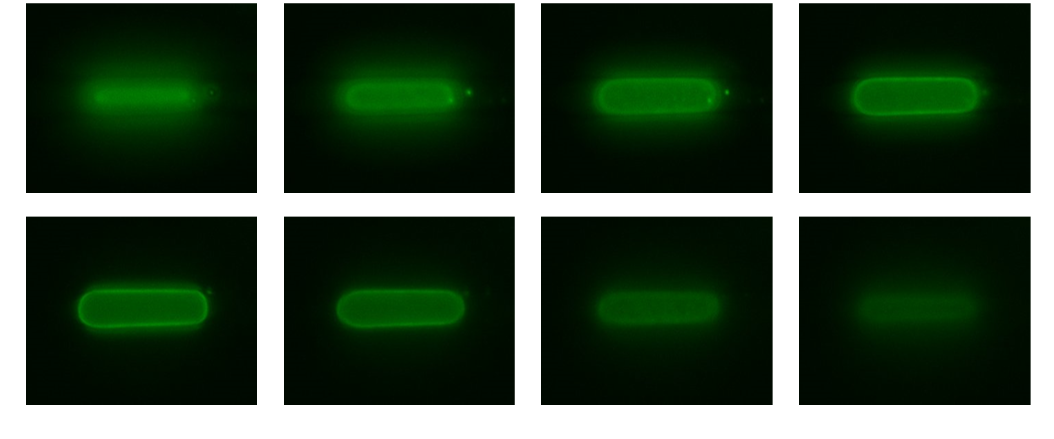

**Supplementary Figure 4.** **Effect of buffer osmolarity on the physical properties and traversal of RBCs.**

Behaviour of RBCs subjected to different buffer osmolarities. (a,b) Mean values and first and third quartile values for the volume and surface area for RBCs subjected to different osmolarities. (c) An inverse linear relationship between MCD and SA:V ratio.

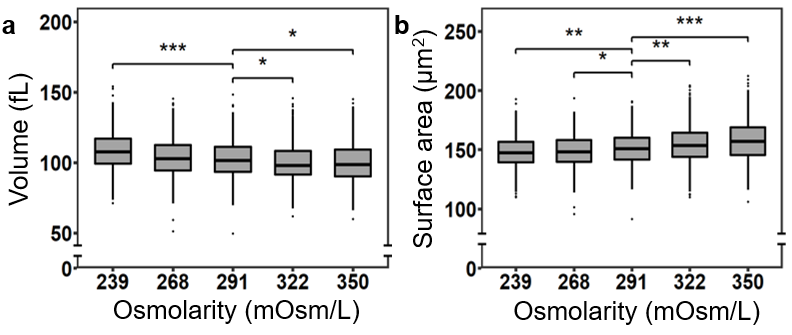

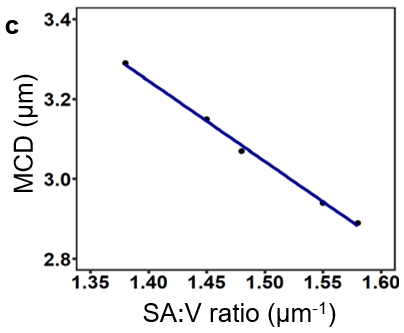

**Supplementary Figure 5. Traversal of *P. falciparum* (CS2) trophozoite-infected RBCs into microchannels.**

Comparison of (a) SA:V ratio and (b) minimum cylindrical diameter (MCD) in uninfected RBCs (blue) and CS2 strain trophozoite-infected RBCs (22-26 hours post invasion) (red).

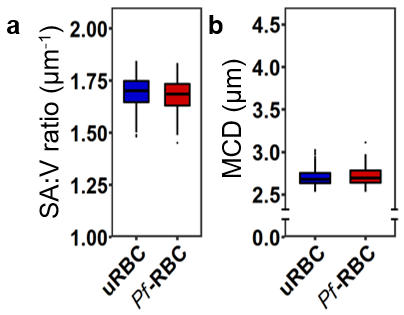

**Figure S6. Reticulocytes are more crenulated than mature RBCs.**

Mature RBC (a) and reticulocyte (b) 3D models derived from block-face SEM serial sections of fixed, dehydrated, resin embedded samples. Remnants of cellular components are visible inside the reticulocyte (c, blue), and can be observed inside cellular blebs (arrows). Scale bars are 1 μm. See Supplementary Video 1 for rotations of the reticulocyte models.

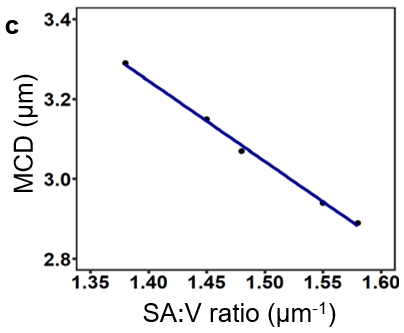

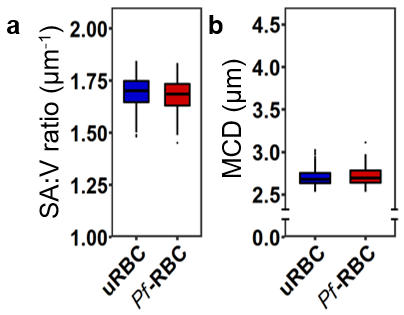

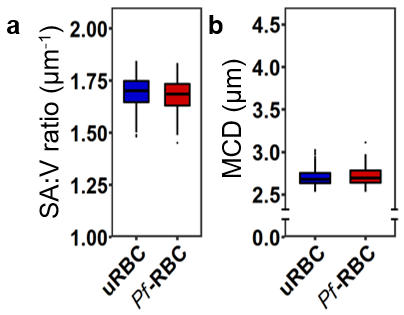

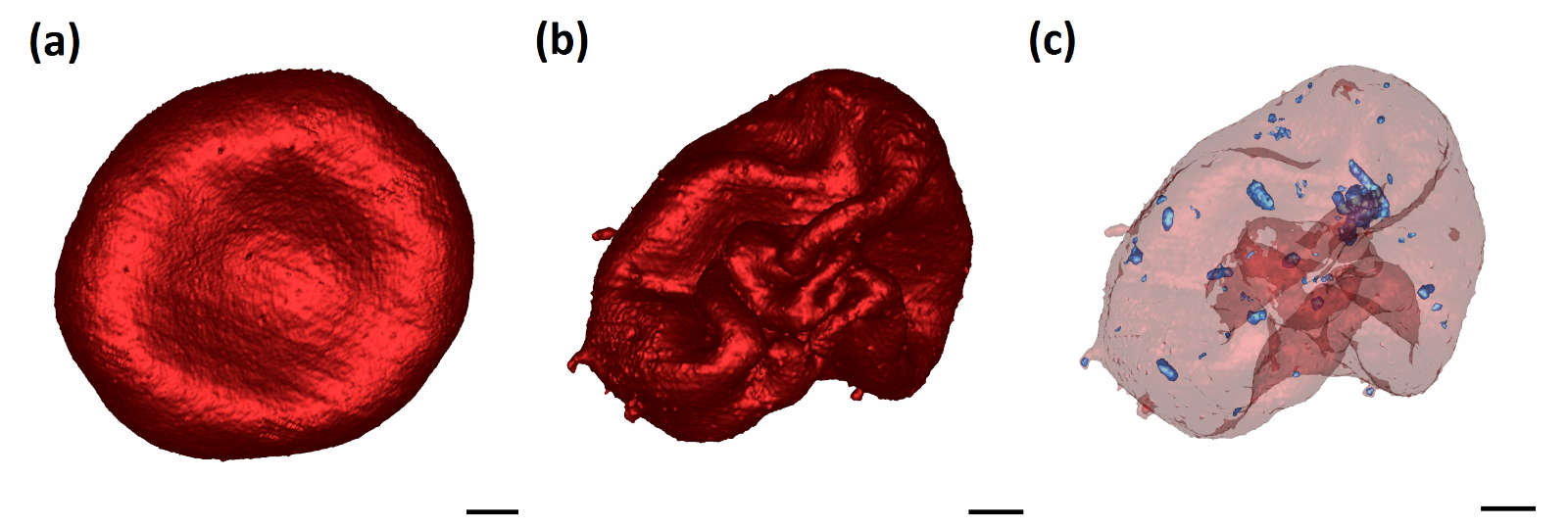

**Supplementary Figure 7. Verifying the 3D** **Finite Element (FE) model with experimental data.**

A pressure analysis was performed to find the best-fit value for the pressure drop to which RBCs are exposed to in the microchannels. Using a SA:V ratio of 1.42 and a shear modulus of 7.3 μN/m, we increased the pressure drop gradually to 4.4 Pa and determined the minimum cylindrical diameter and the length of the elongated RBC. (a) The minimum cylindrical diameter that a normal RBC reaches as a function of the applied pressure drop. (b) The elongated length of the RBC as a function of the applied pressure drop. The green zone shows experimental data for RBCs with similar volume and surface area. Applying a pressure drop of 2.2 Pa to the RBC permits good agreement with the experimental data obtained from 33 cells with similar volume and surface area and a SA:V ratio of 1.42.

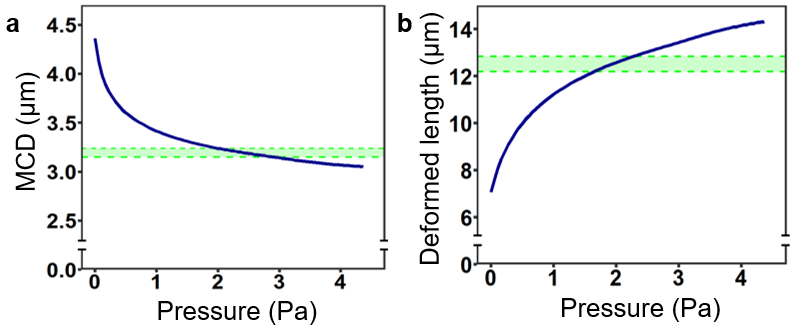

**Supplementary Figure 8. Strain and stress in RBCs subjected to different shear moduli.**

Strain and stress in RBCs with shear moduli ranging from 7.3-109.5 μN/m. (a) RBCs have a highly elastic membrane, which enables them to undergo high deformations. Nominal stress-strain curves for all shear moduli have a relatively gentle slope in the strain range of 0-0.6, similar to the levels of strain that RBCs experience in the wedge-shaped microchannels, indicating a high level of flexibility of the RBC membrane. We plotted the Abaqus-reported maximum in-plane principal logarithmic strain and von Mises stress at the integration point of all membrane shell elements. (b,c) Ranges of strain and stress were calculated at the position of 60 μm inside the microchannels for cell stiffness ranging from 7.3-109.5 μN/m. Our simulations indicate that RBCs exhibiting a shear modulus less than 58.4 μN/mexperience similar strain to each other; however, both strain and stress increased substantially as the shear modulus increased to more than 73 μN/m. Formation of creases within the membrane was the main cause of the increase in strain and stress in such rigid cells. (d,e). We calculated the range of maximum in-plane principal logarithmic strain and von Mises stress in RBCs with various SA:V ratio ranging from 1.15-1.76. Our simulations indicate that RBCs experience a higher range of strain and stress as the SA:V ratio is reduced. This is, the RBC morphology approaches a sphere as the SA:V ratio decreases, increasing the strain associated with squeezing into the narrow microchannels.

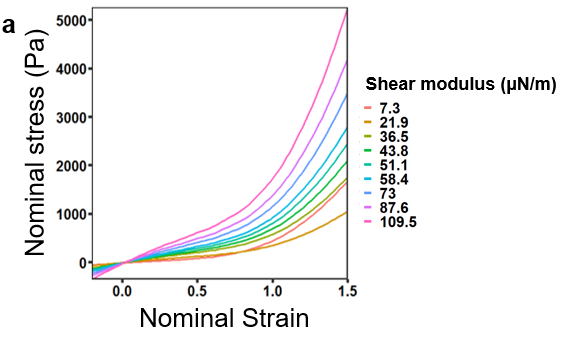

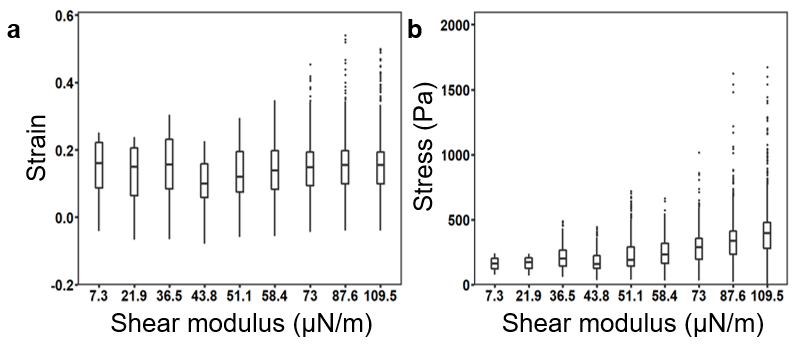

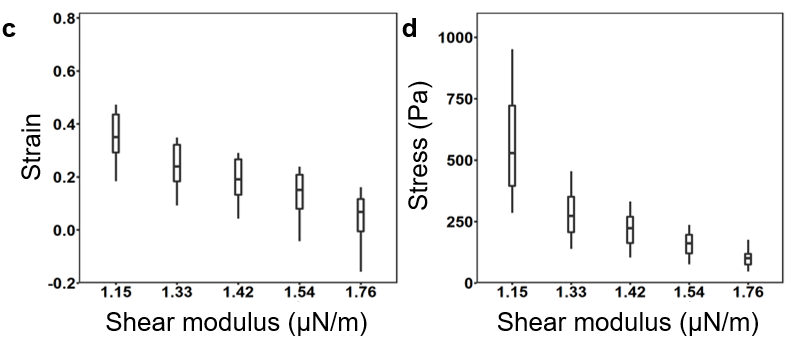

**Supplementary Figure 9. Numerical simulations of the passage of RBCs with a SA:V ratio of 1.42 through microchannels and capillaries as the membrane shear modulus increases.**

Numerical simulations of the passage of RBCs with a SA:V ratio of 1.42 through microchannels and model capillaries as the membrane shear modulus increases. (a) The MCD that RBCs traverse in the microchannels. (b) The minimum required pressure for the passage of RBCs through small cerebral capillaries.

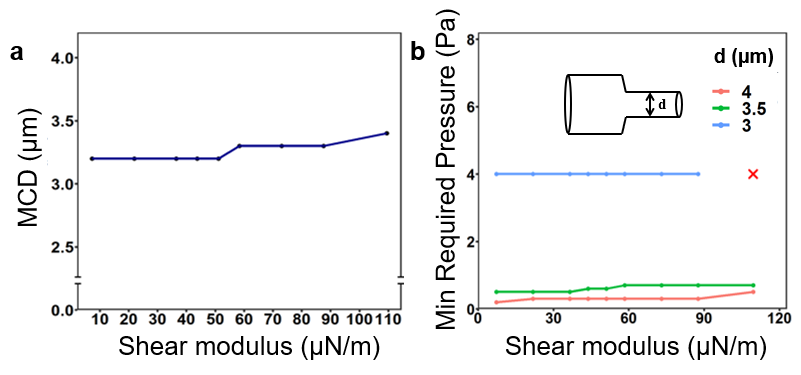

**Supplementary Video 1.** Rotating models constructed from SBF-SEM of an immature reticulocyte. 3D models were derived from block-face SEM serial sections of fixed, dehydrated, resin embedded samples. The surface is rendered solid (left) or translucent (right) revealing remnants of cellular components (blue), and cellular blebs at the surface. Scale bars are 1 μm.
